# Diffusion within the synaptonemal complex can account for signal transduction along meiotic chromosomes

**DOI:** 10.1101/2024.05.22.595404

**Authors:** Lexy von Diezmann, Chloe Bristow, Ofer Rog

## Abstract

Meiotic chromosomes efficiently transduce information along their length to regulate the distribution of genetic exchanges within and between chromosomes. However, the mode of signal transduction remains unknown. Recently, a conserved chromosomal interface, the synaptonemal complex, was shown to be a biomolecular condensate, offering an attractive mechanism for signal transduction: diffusion of signaling molecules within the synaptonemal complex to allow transmission of information along each pair of chromosomes. Here, we test the feasibility of this mechanism in live *C. elegans* gonads. Single-molecule tracking shows that a component of the synaptonemal complex (SYP-3) and a conserved regulator of exchanges (ZHP-3) both diffuse within the synaptonemal complex. However, ZHP-3 diffuses 4- and 9-fold faster than SYP-3 before and after crossovers formation, respectively. We use these measurements to parameterize a physical model for signal transduction. We find that ZHP-3, but not SYP-3, explores the lengths of chromosomes on the time scale of crossover maturation, consistent with a role in the spatial regulation of exchanges. Given the conservation of ZHP-3 paralogs across eukaryotes, we propose that diffusion within the synaptonemal complex may be a conserved mechanism of meiotic regulation. More broadly, our work explores how diffusion contained within condensates regulates crucial cellular functions.

## Introduction

During meiotic prophase, parental chromosomes align with one another, receive double-strand breaks, and select a subset of these breaks to be repaired as physical exchanges known as crossovers. This process is tightly regulated to restrict excessive crossovers while ensuring at least one crossover on each chromosome^1,2^. Central to crossover regulation is patterning by ’crossover interference’: when more than one crossover form on the same chromosome, they are spaced farther apart than would be expected by chance^3^. This suggests that an inhibitory signal propagates along the lengths of paired chromosomes, but the identity of this signal and its mode of transduction are unknown.

The synaptonemal complex has been identified as a major regulator of crossover patterning. It comprises a linear axis parallel to the chromosomes made up of chromatin-associated axis proteins from the HORMA domain family (in worms, HTP-1/2/3 and HIM-3), and a central region (SC-CR) composed of proteins that span the inter-chromosomal space (in worms, SYP- 1/2/3/4/5/6 and SKR-1/2^4,5^). In addition to the intimate association between the parental chromosomes, the synaptonemal complex directly regulates crossovers: multiple proteins required for crossover formation are recruited by the SC-CR, and perturbation of the SC-CR alters crossover interference^6–9^. Recently, FRAP and photoconversion microscopy in *C. elegans* showed that SC-CR subunits relocalize on the minutes scale^10–12^, indicating the SC-CR has liquid properties and suggesting it is a biomolecular condensate.

While many mechanisms have been proposed to implement the chromosomal communication that underlies crossover interference, here we focus on two broad families of models^13,14^. In one set of models, a property of the chromosomal scaffold itself serves as the crossover interference ’signal’. In these models, crossover formation effects a large-scale change to the SC-CR and/or chromatin, which regionally inhibits crossover formation. Such a regional change has been suggested to be release of mechanical strain^15^, although it could occur via a phase change, e.g., in the physical properties of a liquid SC-CR. In second family of models, the diffusion of pro-crossover proteins along the SC-CR, and their subsequent capture at emerging crossovers, is what regulates crossover number. This model is supported not only by the liquid properties of the SC-CR, but also by the localization patterns of pro-crossover signaling proteins. In particular, members of the Zip3 protein family in worms, flies and plants transition from a uniform distribution along the SC-CR to concentrated foci at the repair sites that will become crossovers^16–18^. A hitherto untested prediction of this model is that pro-crossover molecules must be able to diffuse along the SC-CR during the crossover designation stage in order to efficiently sample the range over which interference acts, which is, in worms, the entire length of the chromosome^13,14^

Here, we directly test the feasibility of diffusion-based models to communicate along the chromosome on physiologically relevant timescales. We use single-molecule tracking, super-resolution imaging, and photoconversion microscopy of synaptonemal complex scaffold and client molecules in meiotic nuclei of the *C. elegans* gonad. We establish the extent of these molecules’ motion and changes in motion following crossover formation. By using these empirical parameters in a physical model of protein relocalization, we demonstrate that the behavior of SC-CR clients, but not SC-CR scaffold proteins, is consistent with diffusion-based models of crossover regulation.

## Results and Discussion

### Tracking single molecules on meiotic chromosomes

To understand signal transduction along meiotic chromosomes, we tracked single molecules in living *C. elegans* gonads. We used our recently developed technique^19^, but adapted the dye and imaging modality in a crucial way. Previously, we used meiotic proteins covalently attached to the photoactivatable organic dye PAJF549 using the Halo label^20,21^, allowing sparse photoactivation followed by tracking individual proteins until photobleaching (∼1 to 20 seconds at 10 Hz). However, a large fraction of the tracks appeared within the gonad syncytium, suggesting relatively low penetrance of this dye into nuclei (Fig. S5). While assessing alternative labeling strategies, we serendipitously found that the more permeant dye JF552^22^ could be shifted into a long-lived dark state following high-intensity 561 nm excitation (“bleachdown”).

While most molecules are permanently photobleached by this approach, a sizeable fraction stochastically reactivated over time or when exposed to violet (405 nm) or blue (488 nm) light, analogous to the behavior recently reported for JFX650^23^. By titrating dye concentrations and the extent of the initial bleachdown, we were able to maintain single-molecule concentrations and found that most trajectories were within nuclei (Fig. S5).

We focused on three proteins that represent key elements in meiotic chromosomes: HTP-3, SYP-3, and ZHP-3 (Fig 1A). HTP-3 is a HORMA-domain-containing component of the meiotic axis. To track it we used a mostly functional *htp-3-halo* transgene that we recently generated and characterized^19^. SYP-3 is a component of the SC-CR that was previously tagged at its N-terminus^24^. We tracked it using a *halo-syp-3*, a functional fusion we generated at the endogenous locus and that exhibited fertility indistinguishable from wild-type worms (Fig. S1).

**Figure 1.**
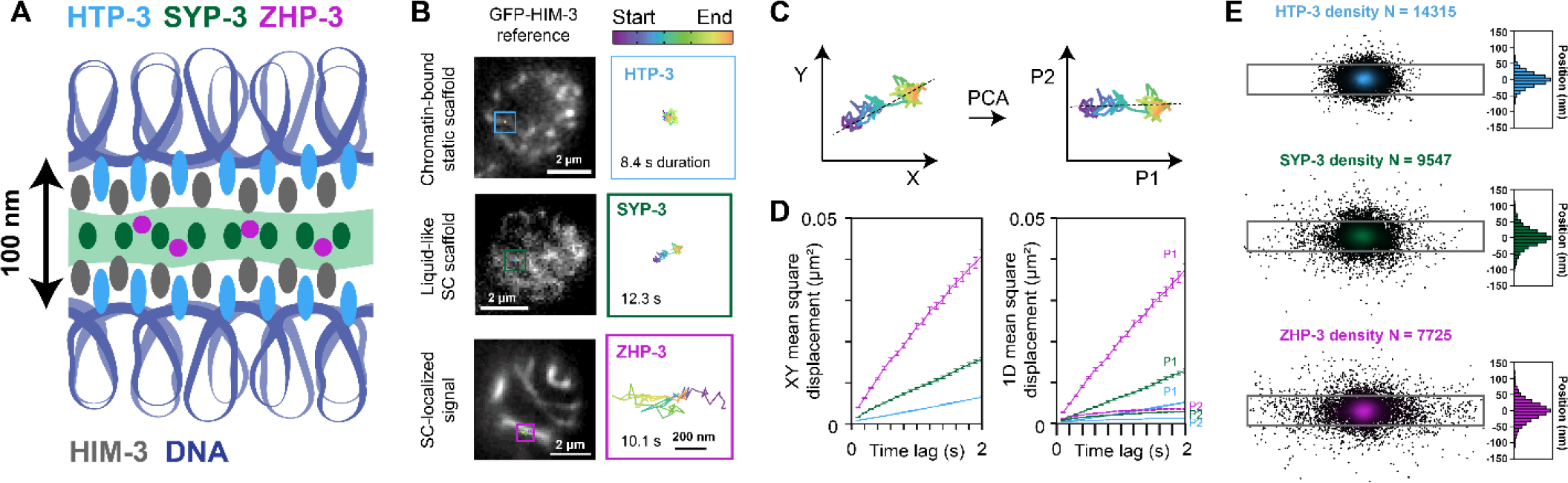
Synaptonemal complex scaffold and client molecules diffuse locally in one dimension A. The organization of proteins spanning the interchromosome region during meiotic prophase I in *C. elegans*. B. Representative tracks of HTP-3, SYP-3, and ZHP-3 (rainbow colored) relative to GFP-HIM-3 (grayscale) in live nuclei of the *C. elegans* gonad. C. Principal component analysis (PCA) identifies the main axis of diffusion for chromosomally-associated tracks. D. Mean square displacement analysis of HTP-3, SYP-3, and ZHP-3. In the right panel the two major PCA components are separated, demonstrating that one-dimensional motion on the P1 axis describes most protein motion. E. Cumulative localizations from all trajectories of each protein, color-coded as a function of local density.

Obtaining a functionally tagged version of the crossover regulator ZHP-3 proved more challenging. ZHP-3 is a member of the Zip3 family of pro-crossover signaling proteins, which contain a RING domain at the N terminus and a large serine-rich region at the C-terminus^17^. Because the RING domain is likely to be functionally critical, we tagged *zhp-3* with *halo* at its C-terminus, but this also detrimentally affected meiotic functions. *zhp-3-halo* homozygous worms had almost no viable progeny, and an average of only four out of the six chromosomes formed chiasmata (Fig. S1). Notably, ZHP-3 localization to meiotic chromosomes was unaffected: like wild-type ZHP-3, ZHP-3-Halo localized to the SC-CR in mid-pachytene before re-localizing to SC-CR-associated foci. The number of foci (4.3 ± 0.9) matched the number of chiasmata (4.1 ± 1.3), suggesting that the tagged version can promote crossover formation, albeit in reduced efficiency relative to untagged ZHP-3. When heterozygous, *zhp-3-halo* did not have any discernible effects on meiosis, and colocalized with a functional allele, *zhp-3-flag* (ref. ^16^ and Fig. S1 and S2). Conducting experiments on both homozygous and heterozygous worms, we found that the dynamics of ZHP-3 were similar overall (Fig. S3). For comparisons with Halo-SYP-3 and HTP-3-Halo, below, we tracked ZHP-3-Halo in heterozygous animals, where meiotic progression was minimally affected.

We obtained single-molecule traces from worm strains harboring Halo-labeled proteins covalently attached to JF552. To place these trajectories into the context of the SC-CR, we recorded diffraction-limited widefield images of the chromosome axis protein GFP-HIM-3 at 30 second intervals (Fig 1B). We began by analyzing diffusion in early pachytene, before crossovers form^10^. We observed distinct diffusive populations: a fast-moving population that rapidly explored the nucleoplasm (which we previously found^19^ to have a D value ≥ 0.1 µm^2^/s), and a slow-moving population associated with the synaptonemal complex (Fig. 1B).

We then analyzed the motion of HTP-3, SYP-3, and ZHP-3 molecules that were associated with the synaptonemal complex. To ensure sufficient statistics, we used trajectories that were at least 1 s in duration and contained localizations in at least 50% of the frames. The resulting trajectories had a median length of ∼3 seconds (Table 1) with average lateral localization precisions of 14 nm to 24 nm. To evaluate the volume explored by these molecules, we used principal component analysis to align trajectories within a local coordinate system (Fig 1C, methods). Strikingly, 80% or more of these trajectories’ motion was contained within the first principal component, P1 (Fig. 1D). By analyzing the increase in mean square displacement along the P1 axis, we measured one-dimensional diffusivity values of D = 0.9, 3.2, and 11.9 x 10^-3^ µm^2^/s for HTP-3, SYP-3 and ZHP-3, respectively (Table 1). The shape of the MSD plot was close to linear. This suggested that these molecules diffused by Brownian motion rather than directed flow or motor-driven motion (which would have generated upward curves), and that motion was not significantly confined on this timescale (which would have generated downward curves). In the second principal component, P2, motion was minimal and the MSD plot curved downward, consistent with restriction to the 100 nm wide SC-CR (Fig. 1D). To visualize this, we plotted all localizations from trajectories within their shared coordinate system, and indeed found that the P2 component of the trajectories was generally confined to ∼100 nm (Fig. 1E).

**Table 1.** Results from single-molecule trajectory analysis. Statistics of analyzed trajectories ≥ 1.0 s. Measurement error was defined as the standard deviation of D calculated from MSD curves generated from trajectories that were randomly resampled with replacement (i.e., bootstrapping). All diffusivity values are in units of 10^-3^ µm^2^/s. “Early/late” refers to stage in pachytene. “Het/homo” indicates hetero/homozygosity for the *zhp- 3::halo* fusion.

|  | #trajectories | Median duration (s) | D<sub>2D</sub> | D<sub>2D</sub> error | D<sub>1D</sub> | D<sub>1D</sub> error |
| --- | --- | --- | --- | --- | --- | --- |
| HTP-3 early | 365 | 3.2 | 0.68 | 0.04 | 0.95 | 0.07 |
| HTP-3 late | 159 | 3.4 | 0.56 | 0.07 | 0.75 | 0.08 |
| SYP-3 early | 284 | 2.9 | 2.22 | 0.23 | 3.15 | 0.37 |
| SYP-3 late | 505 | 3.6 | 0.69 | 0.07 | 1.00 | 0.12 |
| ZHP-3 het early | 171 | 3.8 | 6.63 | 0.66 | 11.8 | 1.27 |
| ZHP-3 het late | 150 | 3.75 | 5.34 | 0.97 | 9.09 | 1.8 |
| ZHP-3 homo early | 343 | 3.3 | 7.19 | 0.92 | 12.0 | 1.68 |
| ZHP-3 homo late | 353 | 3.6 | 4.64 | 0.46 | 8.14 | 1.08 |

Given that the SC-CR is a biomolecular condensate, the likely interpretation of the 1- dimensional movement we observed is that SYP-3 and ZHP-3 diffuse within and along the SC- CR. For SC-CR components, like SYP-3, this reflects either free motion in a liquid phase, or percolation through vacancies in the SC-CR structure. In the case of ZHP-3, this may entail mixing in the liquid (i.e., competing for the binding sites with the scaffold components) or anchoring to distinct sites on one or more SC-CR subunits. A corollary of the latter idea is that the motion of ZHP-3 might reflect a combination of its own diffusive motion as well as the diffusive motion of the SC-CR subunits with which it is associated. Rarely, we observed SYP-3 or ZHP-3 trajectories that switched between motion in the nucleoplasm and motion in the SC (Fig S4), suggesting that the timescale of such association/dissociation events is >10s.

Together, these results indicate that both SYP-3 and ZHP-3 move along the length of chromosomes while remaining mostly constrained to a single SC-CR compartment.

### ZHP-3, but not SYP-3, remains highly diffusive following crossover formation

Fluorescence recovery after photobleaching (FRAP) analysis demonstrated’ that the SC-CR components SYP-2 and SYP-3 become less dynamic as meiotic prophase I progresses, and it has been suggested that this represents a response to crossover formation^11,25,26^. Notably, FRAP reflects the combination of intra-chromosomal diffusion, as measured here, but also binding/unbinding events, which could be inter- and intra-chromosomal. To determine how meiotic progression specifically affects synaptonemal complex-associated diffusion, we compared single-molecule tracking data acquired from nuclei in early and late pachytene (Fig. 2). Averaging many single molecule tracks to approximate the ensemble behavior in each condition, we found that HTP-3, SYP-3, and ZHP-3 exhibited less diffusion following crossover formation (Fig. 2A-B). This slowdown was particularly dramatic for SYP-3, which exhibited diffusion in late pachytene comparable to that of HTP-3 in early pachytene. This indicates that the slowing of SC-CR FRAP rates during meiosis is at least partially caused by slower diffusion along the SC-CR. By contrast, ZHP-3 molecules in late pachytene diffused only slightly more slowly (11.8 ± 1.3 vs. 9.1 ± 1.8 x 10^-3^ µm^2^/s), though exhibited a greater downward curve in their mean square displacements with time. This could indicate confinement along the length of the SC-CR, or might reflect a ’static’ subpopulation of ZHP-3 molecules, potentially associated with repair intermediates.

**Figure 2.**
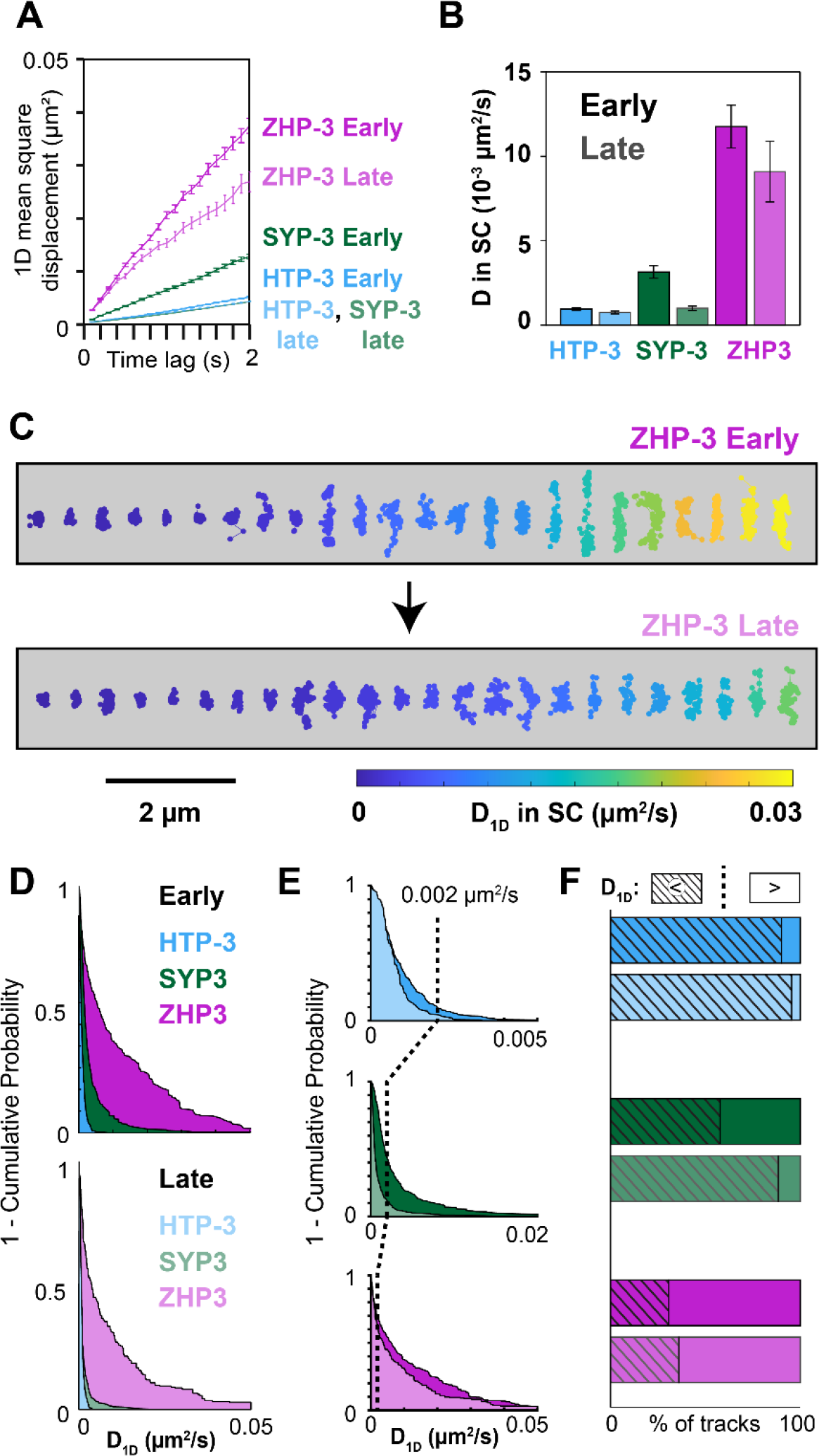
Single-molecule diffusivities shift in response to crossover designation. A. Mean square displacement data along the principal axis of variation. B. Calculated diffusivity from the MSD plots. Errorbars: standard deviation of bootstrapped analyses. C. Example ZHP-3 trajectories before and after crossover formation, aligned vertically and sorted/colored by diffusivity. The longest trajectories from each dataset are shown, and all contain at least 50 localizations. D. Distribution of diffusivities for track populations for HTP-3, SYP-3, and ZHP-3 in early and late pachytene. E. Comparison of early and late pachytene diffusivities for HTP-3, SYP-3, and ZHP-3. The value 0.002 µm^2^/s is indicated in each graph for scale. F. Relative proportion of trajectories below and above the value D1D = 0.002 µm^2^/s for each protein.

To evaluate the underlying distribution of molecular diffusivities, we analyzed diffusion on a per- trajectory basis, and found that motion was highly heterogenous (Fig. 2C). Plotting the cumulative distribution functions of D for these proteins corroborated our results: While a large population of ZHP-3 molecules remained highly mobile after crossover formation, most SYP-3 molecules were not (Fig. 2D-E: HTP-3 exhibited very few trajectories with substantial movement along chromosomes at either stage). Using the 90th percentile of HTP-3 trajectory diffusivity in early pachytene (0.002 µm^2^/s) as a cutoff, we find that 42% and 69% of SYP-3 and ZHP-3 were above that cutoff in early pachytene, versus 12% and 64% in late pachytene, respectively (Fig. 2F).

### Intra-chromosomal diffusion of SC components observed through bulk measurement

To observe intra-chromosomal SC-CR diffusion through an orthogonal approach, we performed high-resolution photo-conversion experiments using confocal microscopy (Methods). We used strains where SYP-3 and the HTP-3-interacting axis component HIM-3 were tagged with the green-to-red fluorescent protein mMaple3^10^. We minimized optical aberrations and background autofluorescence by analyzing dissected gonads rather than the whole worms used in previous efforts^10–12^.

We photo-converted molecules on one segment of a single chromosome (time 0) and took z-stacks every 5 minutes. We traced the chromosome in 3D and measured the length of the activated region (Fig. 3). In line with previous observations^10^, HIM-3 minimally spread along the chromosome in both early- and late-pachytene, exhibiting rates of 10 and 12 nm per minute, respectively. SYP-3, however, spread along early-pachytene chromosomes at an average rate of 260 nm per minute. Consistent with slower fluorescence recovery measured for SYP-3^11^, late-pachytene nuclei exhibited much slower rate of spread, 13 nm per minute, similar to HIM-3 (Fig. 3). This rate translates to spread along the ∼6 μm meiotic chromosomes within ∼23 minutes for SYP-3 in early pachytene and 7-10 hours for HIM-3 and for SYP-3 in late pachytene.

**Figure 3:**
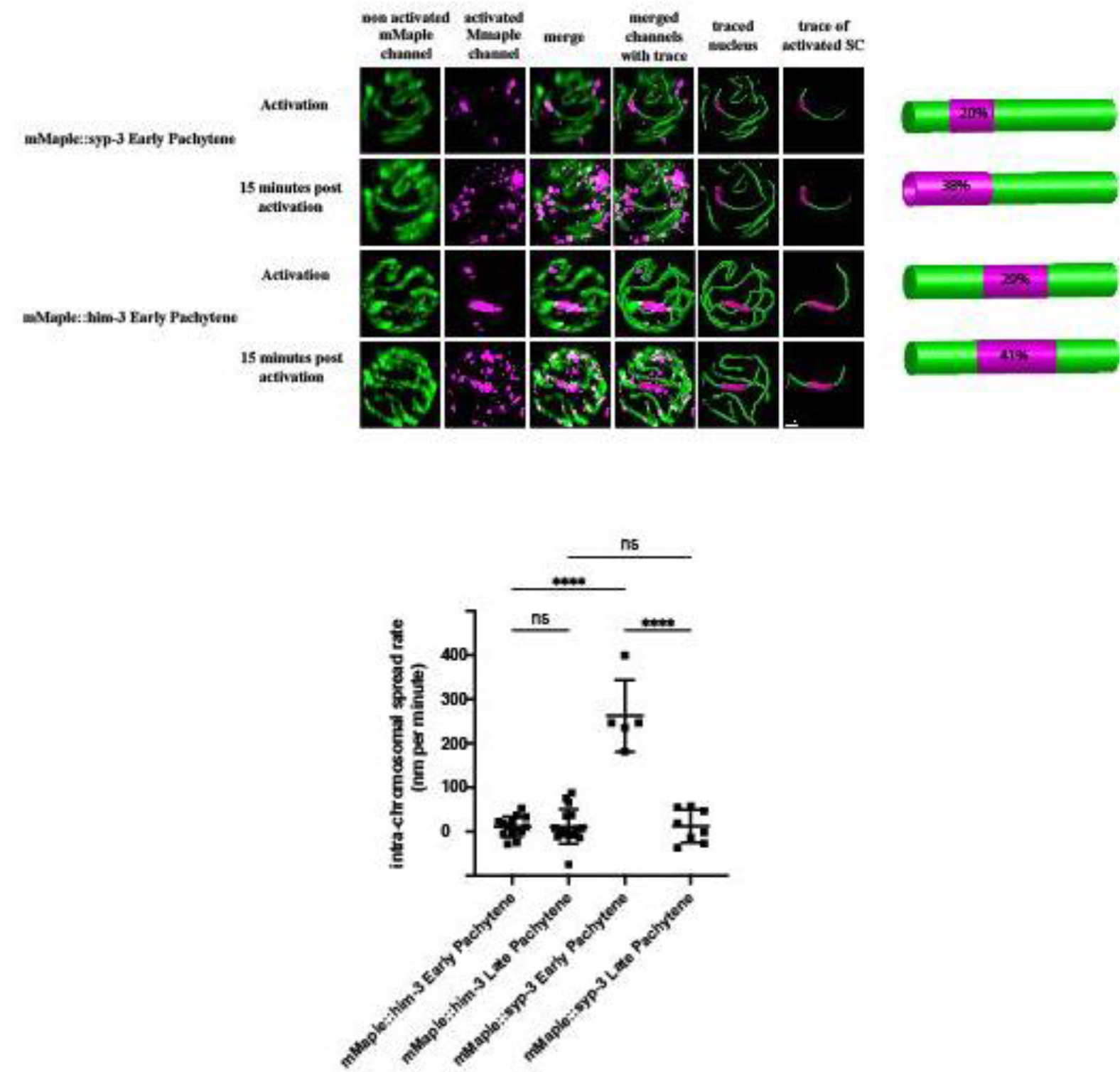
Photoconversion measurement of intra-chromosomal diffusion A. Representative nuclei from photoactivation experiments. B. Measured velocities of intra-chromosomal spread.

This analysis confirms that the movement of SC-CR components along the chromosome in early pachytene, measured above through single-molecule tracking, results in discernible bulk reorganization of the SC-CR along a chromosome. Furthermore, the lack of bulk intra-chromosomal reorganization in late pachytene suggests a greater role for inter-chromosomal turnover of SC-CR components, as one-dimensional diffusion appeared largely restricted during that stage of meiosis (Fig. 2).

### Diffusion-and-capture of ZHP-3 could promote rapid designation of crossovers

The key regulatory events accounting for crossover formation seem to occur over tens of minutes to one or two hours. This estimate is based on the appearance and disappearance of foci of repair intermediates and crossovers in worms^27^ and on physical assays of repair intermediates in budding yeast^28,29^. If crossover interference is coordinated by intra-chromosomal diffusion, such diffusion must be able to occur faster than that timescale. For *C. elegans*, which forms one crossover on each chromosome pair ^30^, we expect this requirement to be particularly strict: that is, signals must be transduced along the length of the entire chromosome (∼6 µm) within tens of minutes.

To evaluate the limits of diffusive information flow along the SC-CR, we used our measurements of SYP-3 and ZHP-3 to parametrize simulations of one-dimensional Brownian diffusion (Methods). We examined two kinds of signal transport in which diffusion is the rate-limiting step (Fig. 4). In one, we simulated how molecules emanating from one end of the chromosome equilibrate in concentration on the other end. This approximates the spread of an inhibitory signal from a crossover site to the distal end of the chromosome. In another, we measured the time to ‘trap’ molecules at a sink, representing the depletion of a pro-crossover factor by a repair intermediate. We focused on early pachytene measurements, since at late pachytene, when ZHP-3 has accumulated at crossover foci, designation is thought to have already occurred.

**Figure 4.**
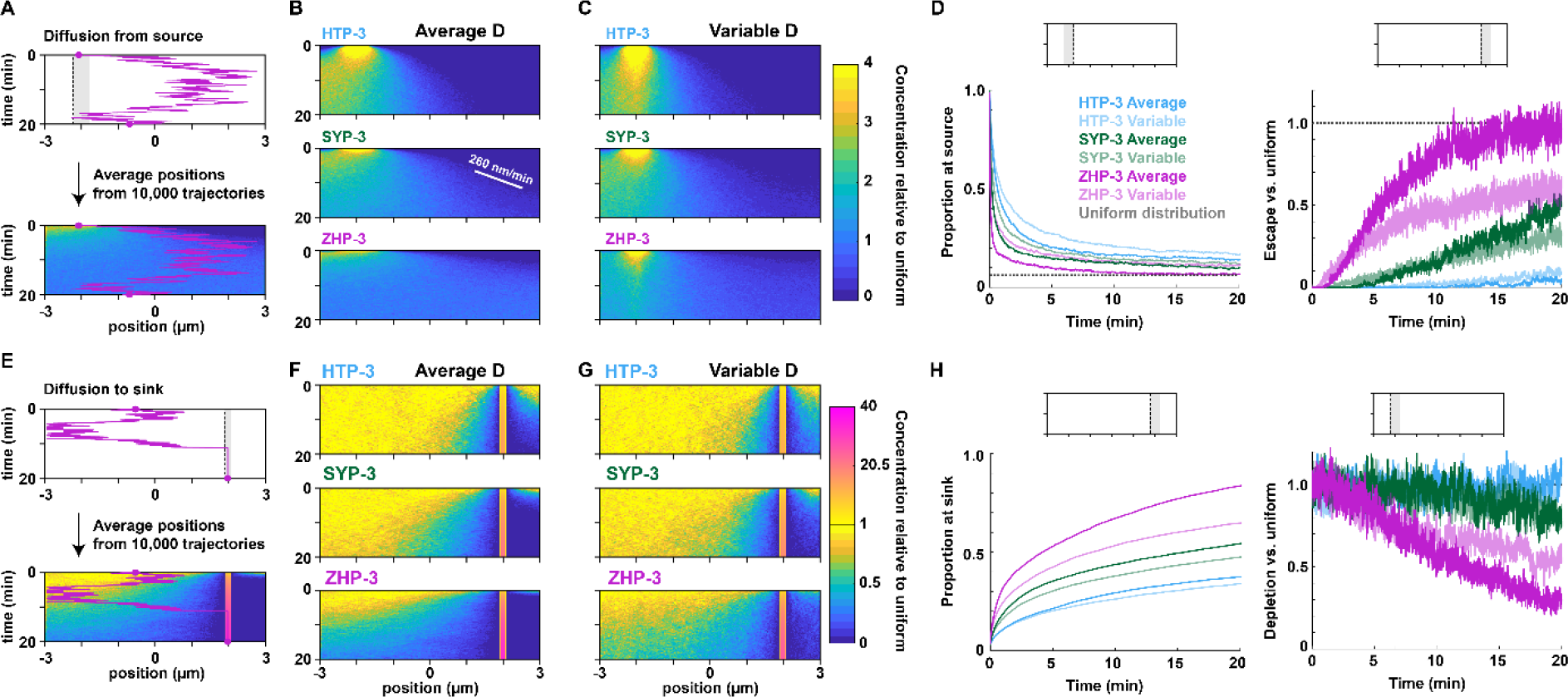
Simulations of relocalization within the synaptonemal complex. A. Example simulated trajectories diffusing from an initial 300 nm wide “source” position. (Empirical diffusivity of ZHP-3.) B. Kymographs of simulated trajectories beginning from a source position using averaged empirical D for each protein. C. Kymographs using variable D drawn from the empirical distribution of each protein. D. Change in number of simulated molecules remaining at the source position (left) or appearing at the distal side of the chromosome (right) relative to a uniform density (dotted line). E. Example simulated trajectories starting from random initial positions and binding irreversibly when reaching a 150 nm wide “sink” position. F. Kymographs of simulated trajectories binding at a sink position using averaged empirical D for each protein. G. Kymographs using variable D drawn from the empirical distribution of each protein. H. Change in number of simulated molecules reaching the “sink” location as a function of time (left) and depleted at the distal side of the chromosome (right).

Simulating the equilibration of HTP-3, SYP-3, and ZHP-3 from a small region 1 µm along a 6 µm chromosome (Fig. 3A-D), we found that ZHP-3 rapidly (∼10 mins) achieved a mostly uniform distribution along the chromosome, while SYP-3 and HTP-3 redistributed approximately 4-fold and 10-fold more slowly, consistent with the inverse relationship between diffusion coefficient and time to diffuse a given distance. Notably, the rate of SYP-3 redistribution was comparable to the velocity measured from our photoconversion experiments of 260 nm/minute (Fig. 3B and white line in Fig. 4B). HTP-3 moved more quickly than observed in our photoconversion measurements of the HORMA domain protein HIM-3. This may reflect either a true difference between these proteins, an overestimate of D due to nuclear or chromosomal motion, or different signal-to-noise threshold in simulation *versus* fluorescence microscopy. We performed these simulations both using the diffusivity calculated from the full molecule population (“Average D”; Fig. 2B) and by assigning a different D value for each simulated trajectory, pulling from the empirical distribution (“Variable D”; Fig. 2D). The simulations maintained similar trends overall, although all exhibited slower overall redistribution in the “variable” case, likely due to the subpopulation of slow-moving molecules remaining at the source position.

We then simulated the rate at which an initially uniform distribution of HTP-3, SYP-3, and ZHP-3 relocalized to a sink 1 µm from the end of the 6 µm chromosome. (Fig. 3E-H). In coarsening models of interference, the timescale reflects both diffusion and the kinetics of trapping at repair sites^14^. By simulating a single sink that instantaneously and irreversibly binds any molecule that reaches it, our simulation reflects the limits of diffusion alone. As expected, all proteins exhibited roughly logarithmic accumulation at crossover sites due to the depletion of molecules from the rest of the chromosome. Depletion occurred most rapidly in the ∼150 nm around the sink, as well as in the short chromosome arm near it. Interestingly, we found that ZHP-3 was capable of 50% depletion at the distal end of the chromosome by 10 minutes and 70% depletion by 20 minutes. By contrast, neither SYP-3 nor HTP-3 were depleted more than 20% by this time.

In sum, our simulations confirm that the dynamic relocalization of ZHP-3 is a particularly good match for the timescale of crossover designation in the coarsening model, with HTP-3 and SYP- 3 making much poorer candidates. In addition, the rapid and efficient depletion of even virtually static components within ∼150 nm of the “sink” suggests that the peri-crossover region may have distinct properties. This could play a role in regulating the local remodeling of the chromosome in response to crossover formation, as has been observed in worms^6,27^. It may also underlie other forms of meiotic regulation, such as the local spatial regulation of double-strand breaks in yeast^31,32^.

### Evidence for trapping of crossover regulators through local aggregation

ZHP-3 and other crossover regulators, like HEI10 in plants^18,33^ and Vilya in flies^17^, accumulate at sites of crossovers as they are depleted from the rest of the chromosome. This accumulation is consistent with reaction-diffusion-type models for crossover regulation. Such models rely on two key principles: the first is a diffusing regulator that samples the chromosomal region under regulation; in the case of worms, that would be the entire chromosome. Here we have demonstrated this ability for ZHP-3; while both SYP-3 and ZHP-3 diffuse inside the synaptonemal complex (Figs. 1 and 2), only ZHP-3 was able to effectively sample the entire 6 µm chromosome in biologically-relevant timescales (Fig. 4). The second principle is competition between exchange precursors for a limited amount of regulator. In particular, such competition must favor the formation of a few (ideally, one) densely concentrated foci at the range over which interference acts.

Such dense accumulation could be achieved through a transition between diffusive association of molecules within the SC-CR and mostly irreversible aggregation at crossover precursors.

Several lines of evidence support this idea. First, 1,6-hexanediol, which dissolves both the SC- CR itself and SC-CR-associated ZHP-3, does not dissolve ZHP-3 foci^10^. Second, different types of cytological preparation fail to localize crossovers inside the SC-CR; instead, crossovers appear to the side of the SC-CR^34,35^ or form a bubble that deforms the axes^27^. Third, ZHP-3 and other crossover markers in worms^10,16^ and HEI10 in plants^9,33^ form chromosome-associated foci at recombination intermediates even in the absence of the SC-CR. The ability to aggregate independently of the SC-CR suggests that the foci that form in the context of the SC-CR are not merely the consequence of highly biased diffusion into, but not out of, the vicinity of crossover precursors.

To test whether the two proposed material states of ZHP-3 - diffusive and aggregated - are reflected in its localization vis-à-vis the SC-CR, we localized ZHP-3 relative to the two parallel axes abutting the SC-CR. We used super-resolution stimulated emission-depletion microscopy (STED), which resolves the ∼150nm separated axes^35^. While ZHP-3 molecules along the chromosomes were tightly localized within the axes with a variance of 0.054, ZHP-3 foci that form in late pachytene and are associated with COSA-1 exhibited a variance in position of 0.126 the interaxis distance (p < 0.011; Fig 5). Such wide distribution is expected for foci that are associated with, but not embedded within, the SC-CR, and is in line with our published finding on other crossover markers^35^.

**Figure 5.**
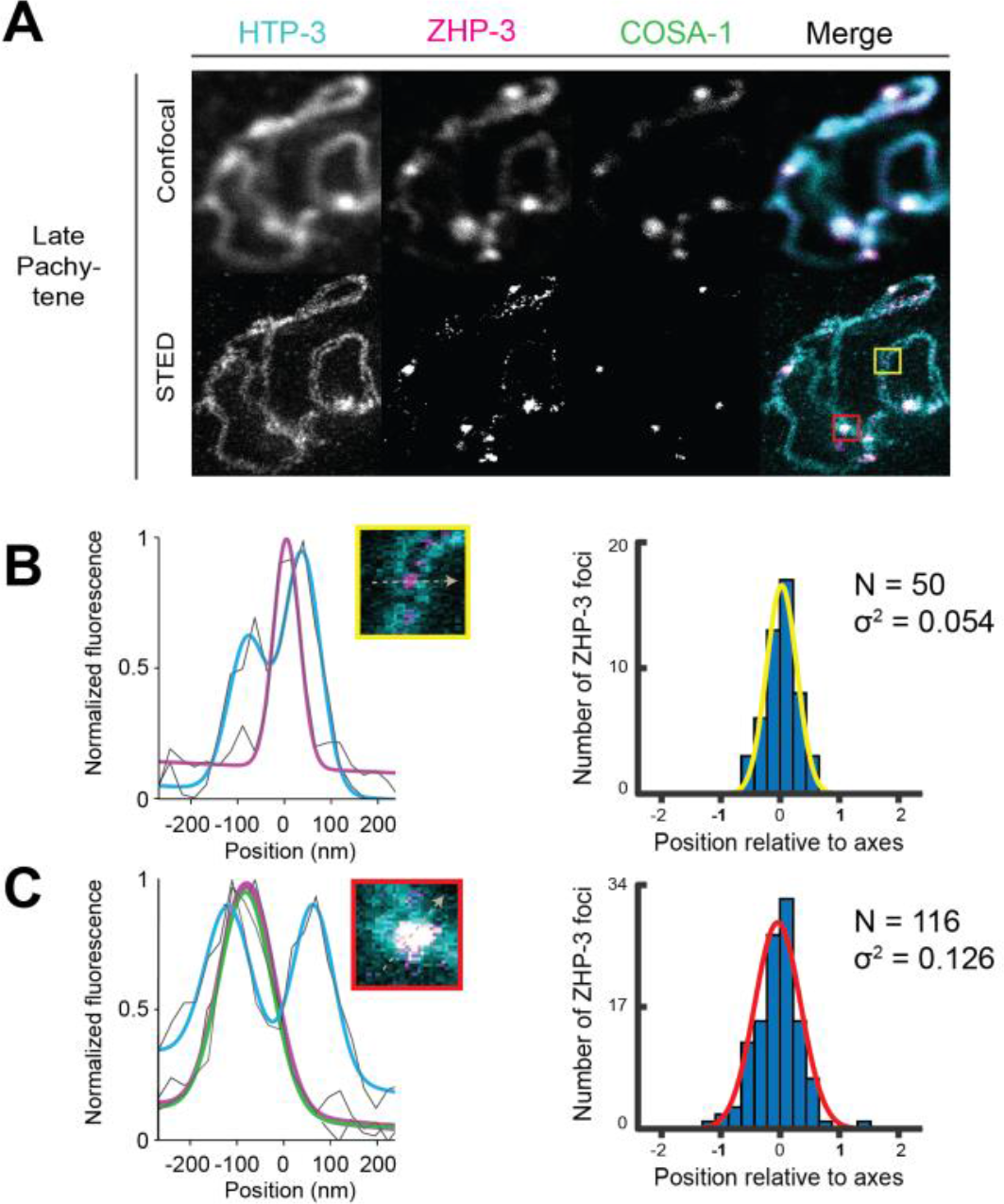
ZHP-3 at crossover sites may not directly associate with the SC. A. Three-color confocal and STED images of a late pachytene nucleus. Example ROIs away from and at crossovers are marked in yellow and red. B. Analysis of the distributions of HTP-3 and ZHP-3 STED data away from crossover sites. Left, fits to line profiles from the yellow ROI, inset. Right, distribution of ZHP-3 positions relative to the axes for many foci. C. Analysis of the distributions of HTP-3, ZHP-3, and COSA-1 STED data at crossover sites. Left, fits to line profiles from the red ROI, inset. Right, distribution of ZHP-3 positions relative to the axes for many foci.

### Implications for crossover regulation on meiotic chromosomes

Here, we showed that SYP-3, a member of the SC-CR scaffold, and ZHP-3, a pro-crossover signal, diffuse along the length of the SC-CR. While both SYP-3 and ZHP-3 slow down with meiotic progression, many ZHP-3 molecules continue to diffuse within the SC-CR. What are the implications of this for crossover regulation? Our results provide a plausible molecular mechanism to implement crossover patterning by coarsening. In this model, ZHP-3 is recruited throughout the chromosome where it can efficiently sample multiple repair intermediates by diffusion and preferentially accumulate at a subset of sites. Accumulation could be driven by spontaneous self-assembly, analogous to Ostwald ripening of phase-separated droplets in solution, or by local stabilization through a biochemical feedback loop. Designated crossovers then act as a sink to deplete SC-CR-associated ZHP-3 and prevent formation of additional sinks, and, consequently, additional crossovers. In our simulations, ZHP-3 diffusion is sufficiently rapid to deplete the SC-CR-associated pool along the length of chromosomes within minutes.

Does confirming the sufficiency of the coarsening model rule out other models based on changes in the SC or chromatin state? Not necessarily. The change of SYP-3 from a more to less diffusive state following crossover formation could change the ability of the SC-CR to remodel as designated crossovers emerge, adding an additional layer of regulation. For instance, the formation of axis “bubbles” (ref. ^27^ and Fig. 5) may require a relatively fluid SC-CR, and slowing of the SC-CR following crossover formation could block crossovers beyond the first. Future efforts to validate coarsening models will likely require additional empirically well-defined parameters. In addition to diffusivity, key parameters of coarsening models^18,36^ include binding/unbinding to SC-CR and to crossover sites, protein copy numbers, and information on post-translational modifications.

An important finding of our work is showing that both scaffold subunits (SYP-3) and recruited client proteins (ZHP-3) can diffuse freely within the SC-CR. Such internal diffusion is a hallmark of liquid organization, indicating that even when assembled between chromosomes, the SC-CR is a liquid. Further, this work establishes that ZHP-3, and potentially other Zip3 family members, are bone fide SC-CR clients. The geometry of meiotic chromosomes may explain the relatively low diffusivity we observed (0.001 to 0.01 µm^2^/s) compared to typical D values found in other condensates (0.01 to 10 µm^2^/s) ^37^. It is likely that the attachment between the axes greatly increases the drag forces on the SC-CR, analogous to the effect of rigid support on the diffusivity of supported lipid bilayers^38^. It is also possible the ordered nature of the SC-CR, with regular 10-15 nm striations observed by electron microscopy^2^, contributes to the slower mobility. However, our general findings of how diffusion sets the rate of signal propagation within the SC- CR are likely to apply more generally, both to other 1D condensates, such as proteins wet to the cytoskeleton^39^ and to transport within multiphase condensates^40^. The unilamellar morphology of the SC-CR makes it an excellent object of *in vivo* imaging. These cytological properties and the distinct outcomes of SC-CR-dependent signaling render the SC-CR an ideal ’model condensate’ in which to empirically probe the functional implications of intra-condensate diffusion.

## Acknowledgements

We thank members of the Rog lab for helpful discussions; Matthew L. Schwartz for providing reagents and guidance related to the SapTrap method; Erik M. Jorgensen and members of the Jorgensen lab for discussion of the research strategy; Jonathan Grimm and Luke Lavis for providing fluorescent dyes used in this work. Worm strains were provided by Abby Dernburg and by the Caenorhabditis Genetics Center, which is funded by NIH Office of Research Infrastructure Programs (P40 OD010440). L.v.D. was supported as a Mark Foundation for Cancer Research Fellow (DRG-2372-19) and is supported as a Dale Frey Scientist (DFS-56-23) by the Damon Runyon Cancer Research Foundation. Work in the Rog lab is funded by R35GM128804 grant from NIGMS.

## Methods

### Worm strains and transgenes

*C. elegans* strains were maintained using standard methods on nematode growth medium agar plates seeded with OP50 *E. coli*. Well-fed worms were maintained at 20°C for at least one generation prior to microscopy, and two generations prior to progeny and male counts. Immunofluorescence, progeny counts, male counts, foci counts and DAPI staining bodies were performed as previously described^7^

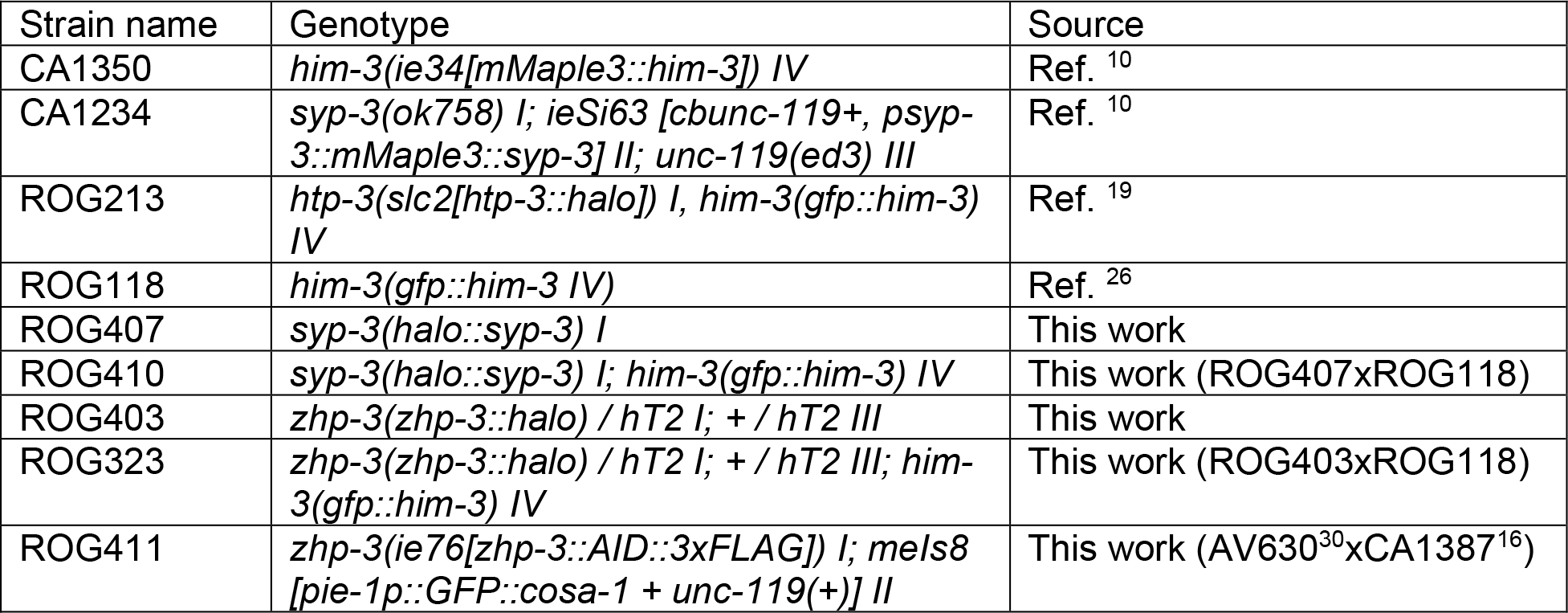

The *halo::syp-3* allele was generated as a CRISPR knock-in at the native *syp-3* site by InVivo Biosystems (Eugene, OR) by direct injection using the sgRNAs ACCGATTCAAAATTTTCTGC<u>GGG</u> and GTTCACCAACTCGCTCGCTG<u>GGG</u>. The homologous template contained 35 base pair homology arms (GTCAGGCAGTAAATGGTGACCGATTCAAAATTTT and CTGGGGAAAGAGGTATGTTTTGAGGCAAACATCAA). The resulting fusion included the first 19 amino acids of SYP-3 (MNFEKLVSQAVNGDRFKIF) followed by a flexible linker (GGTGSGSSTSTG), a synthetic intron, the full 297 amino acid Halo gene, a flexible linker (GGTGSGTGGS) and a synthetic exon coding for the first 19 amino acids of *syp-3* followed by the remainder of the native *syp-3* gene. The progeny counts and percentage of male progeny in strains harboring this transgene were not significantly different from wildtype, indicating it did not perturb the functions of *syp-3* (Fig. S1).

The *zhp-3::halo* allele was generated by injecting plasmids expressing the homology template and sgRNA generated by the SapTrap system^41^ into a strain harboring Cas9 under the *mex-5* promoter and *cre* recombinase under a heat shock promoter^42^, selecting for and sequence- confirming integrants at the *zhp-3* locus. The *unc119* intron wea removed by expressing *cre* via heat shock. ZHP-3 and Halo were connected with the linker AGSGGSGGTGGSGM. The presence or absence of the *unc119* intron did not affect fertility. Due to the low progeny counts for this strain (Fig. S1), strains harboring the *zhp-3::halo* allele were maintained as heterozygotes using the *hT2* balancer chromosome. Experiments with heterozygotes used the balanced strain, while experiments with homozygotes used homozygous progeny of a heterozygous hermaphrodite.

### Single-molecule microscopy

Single-molecule microscopy in *C. elegans* gonad was performed using our previously reported methods^19^ with minor modifications. Young adult hermaphrodites (with a single row of eggs) were dissected in 20-30 µL embryonic culture medium (ECM) without sucrose and with 0.01% Tween 20. The JF552 dye attached to a Halo ligand (JF552-HTL) was added directly to the dissection buffer at a concentration of 0.5-1.0 nM. Agarose pads prepared at a concentration of 2% in water and maintained in a humid chamber with 20 µL ECM added after gelation. Excess liquid was dried in air or by wicking with a Kimwipe immediately prior to depositing dissected worms onto the pad by pipetting. Pads were covered with a 22x22 no. 1.5 glass coverslip, sealed with 1:1:1 Vaseline/Lanolin/Paraffin, and imaged immediately. Data were collected on a biplane single-molecule fluorescence microscope (Vutara 352, Bruker) with a 60x NA 1.3 silicone oil objective (Olympus UPLSAPO60XS2). JF552 was photobleached to reduce the number of active emitters to single-molecule concentration using intense (∼60 μW/μm^2^ at the sample plane) 561 nm light . Because the 488 nm light used to image GFP-HIM-3 reactivated some JF552 molecules, data were acquired in cycles. Each cycle included brief bleachdown illumination (561 nm, ∼60 μW/μm^2^, with intensity progressively lowered in cycles after the first), 30 seconds single-molecule recording (561 nm, ∼4 μW/μm^2^), and one frame of reference data (488 nm, < 1 μW/μm^2^) recorded with 100 ms frames. To avoid artifacts due to time from dissection or variation between gonads, data on early and late pachytene were acquired in the same gonad. For each protein, we acquired at least one pair of datasets in the order “early then late” and one set in the order “late then early.”

### Trajectory analysis

Trajectories were analyzed in a similar way to our previous work^19^. Due to the better precision in XY vs. Z coordinates, only XY information was used in analyzing diffusion. This is also a good approximation of the morphology of the synaptonemal complex, which is mostly flat at the bottom surface of meiotic nuclei, and is roughly parallel to the image plane.

Molecule positions were analyzed from raw data using Vutara SRX software (Bruker). Trajectories were reconstructed from (x, y, z, t) point data using the *swift* software (v. 0.4.3) developed by the Ulrike Endesfelder lab. The same parameters were used for all datasets. Significant parameters included the allowed number of dark frames between localizations (6), the maximum displacement between localizations (400 nm), and the prior probability of bleaching in each frame (0.05), blinking (0.02), and reappearing (0.5). Trajectories were manually inspected to ensure they could be ascribed to single molecules. Frames in which localizations could not be unambiguously assigned (e.g. if bleachdown left too many fluorescent molecules) were excluded from analysis.

We analyzed trajectories that were at least 1 second in duration, contained localizations in at least 50% of frames, and were associated with the SC-CR. We defined synaptonemal complex-associated molecules as those that appeared within ∼300 nm of the GFP-HIM-3 reference image and that did not exhibit rapid nucleoplasmic motion (Fig. S6).

Analysis of trajectories was performed using custom scripts in Matlab (The Mathworks, Inc.; code available upon request). Trajectories were centered by subtracting the mean XY position and aligned by performing principal component analysis (function: *pca*) and rotating data from X-Y to P1-P2 coordinates. Values of the diffusion coefficient D were calculated from linear fits to the mean square displacement data using the equations

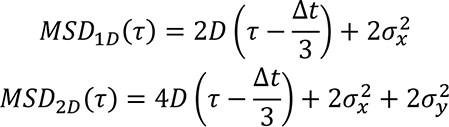

for one-dimensional and two-dimensional diffusion, respectively, time lag τ, exposure time Δ*t*, and localization error σ. Analysis based on MSD1D assumes that the “true” diffusion of molecules is along the one-dimensional SC-CR axis, while analysis based on MSD2D does not require this assumption. Across proteins (HTP-3, SYP-3, ZHP-3) we saw similar trends of diffusivities from each method (Table 1). The precision of these estimates of D was defined as the standard deviation from bootstrapped datasets (i.e., from randomly selected trajectories sampled with replacement). Distributions of D used in downstream analyses were weighted by the number of localizations of each track.

### Simulations

Simulations of molecular diffusion within the SC were generated similar to previous work^13^. One-dimensional displacements were generated from a random distribution with mean 0 and variance 2DΔt for D the diffusion coefficient and Δt the step duration. For these simulations we used Δt = 50 ms. Trajectories were confined to a 6 µm range with absorbing (nonreflective) walls. Kymographs were reconstructed at a sampling rate of 200 ms in time and 20 nm in space. Trajectory start positions were drawn from a uniform distribution in the range [-3, 3] µm. For simulations based on escape from a source position, this range was constrained to [-2.15, -1.85] µm. For simulations based on capture at a sink, trajectories were defined as trapped at a random position within the range [1.925, 2.075] µm upon their first entry into this range.

### Photoconversion microscopy and analysis

Photoconversion microscopy samples were prepared as described for single-molecule microscopy (above). Data was collected with a Zeiss LSM880 confocal microscope with Airyscan in SR mode, equipped with a x63 oil objective. Only fully intact gonads dissected less than 40 minutes prior to imaging were used. Late pachytene was defined as 3 rows prior to the pachytene-to-diplotene transition; early pachytene was defined as 3-4 rows following the zygotene-to-pachytene transition. Diffraction-limited regions were selected for photoconversion by 405 nm light in 4-6 nuclei. Regions were selected such that only one SC-CR stretch would be activated in each nucleus. Prior to photoconversion, the field of view was pre-bleached using 561 nm 75-100% laser power for 30-45 secs going up and down in Z to decrease background noise and bleach pre-activated mMaple3. Following photoconversion, data were collected as z-stacks with laser powers set at 2-2.5% 488 nm and 5.5-7.3% 561 nm, and 1-2 μs pixel dwell times. Data were collected at 5 minute intervals for 30 minutes or until the nuclei drifted out of the imaging volume. Images were Airyscan-processed (ZEN Blue, Zeiss).

Datasets were imported to Imaris (Oxford Instruments) and super-sampled in z to generate cubical voxels. Data were cropped to single nuclei and the length of the activated region was measured at every time point using the Point Measurement tool in manual auto-depth mode. Contrast was altered to minimize background signal and show the full extent of the activated section. Intensity graphs were generated in Imaris using the Point Measurement tool.

### STED microscopy and analysis

We performed STED microscopy and line profile analysis using our previously published approach^35^ with ROG411 worms. Primary antibodies were: mouse anti-FLAG, chicken anti-GFP, and guinea pig anti-HTP-3. Secondary antibodies were: goat anti-mouse STAR 460L (Aberrior), donkey anti-chicken Alexa 594 (Jackson ImmunoResearch), and goat anti-guinea pig STARRED (Aberrior), all at 1:200. Slides were mounted in 12 µL Aberrior Mount liquid antifade.

### Statistical Analysis

Statistical significance between the distributions shown in Figure 5 was calculated using Bartlett’s test. The data in Figure S1 was analyzed in Prism 10 (GraphPad). * indicates p<0.05; ** indicates p<0.005; *** p<0.0005; **** p<0.0001. Only statistically significant comparisons are shown.

## Supplemental Data

**Figure S1:**
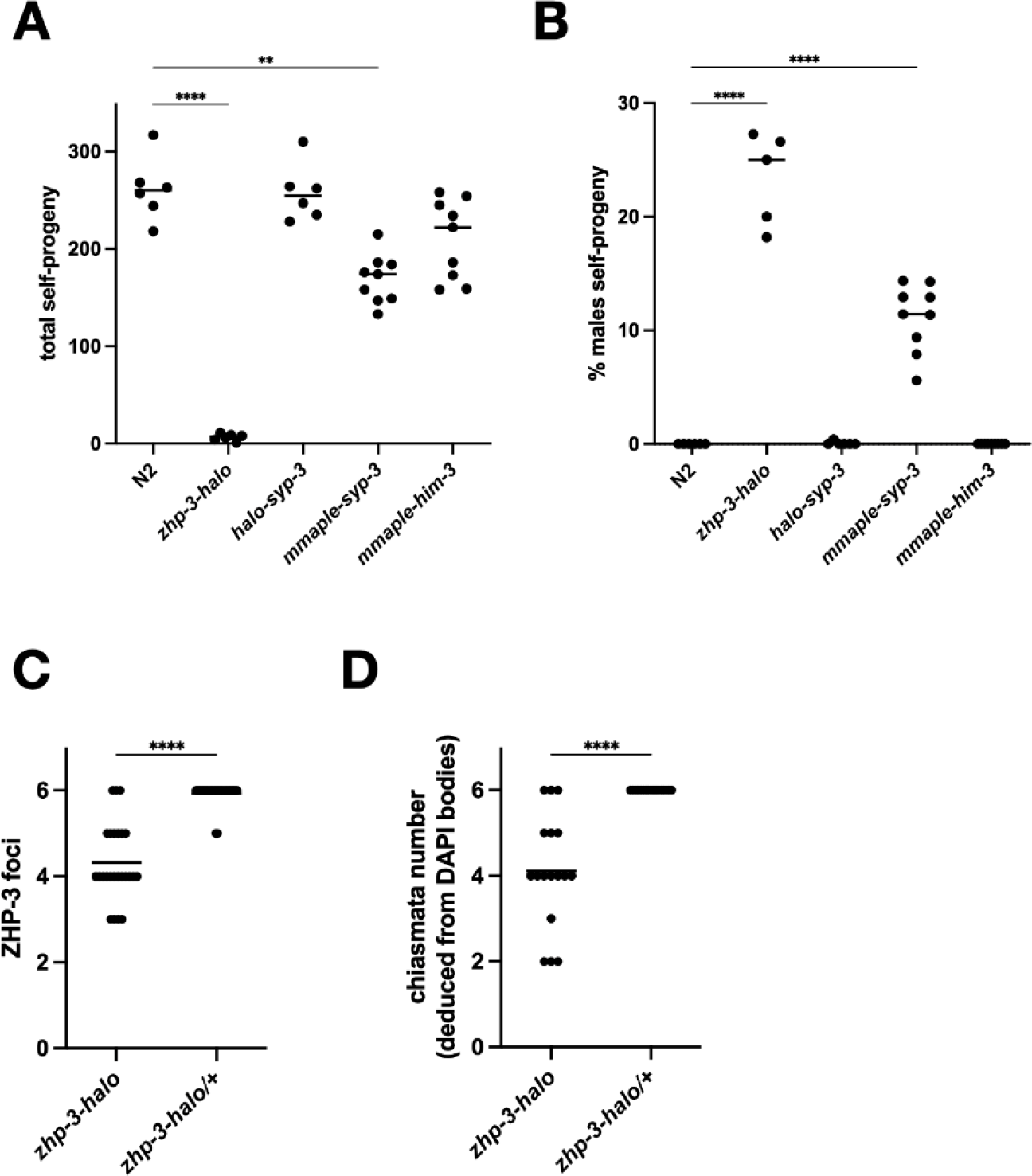
Functional characterization of transgenes used in this study A. Total self-progeny from adult hermaphrodites of the indicated genotypes. Each data point indicates the complete progeny of a single worm. B. Percent male self-progeny from complete broods of worms of the indicated genotype. C-D. Meiosis is minimally perturbed in *zhp-3-halo/+* worms. C. Number of ZHP-3 foci per nucleus. Wild-type worms have 6 foci, one per each pair of homologous chromosomes. D. Number of chiasmata deduces from the number of chromosomes in diakinesis (“DAPI bodies”). Worms have 12 chromosomes, with each additional chiasmata reducing the number of DAPI bodies by one.

**Figure S2:**
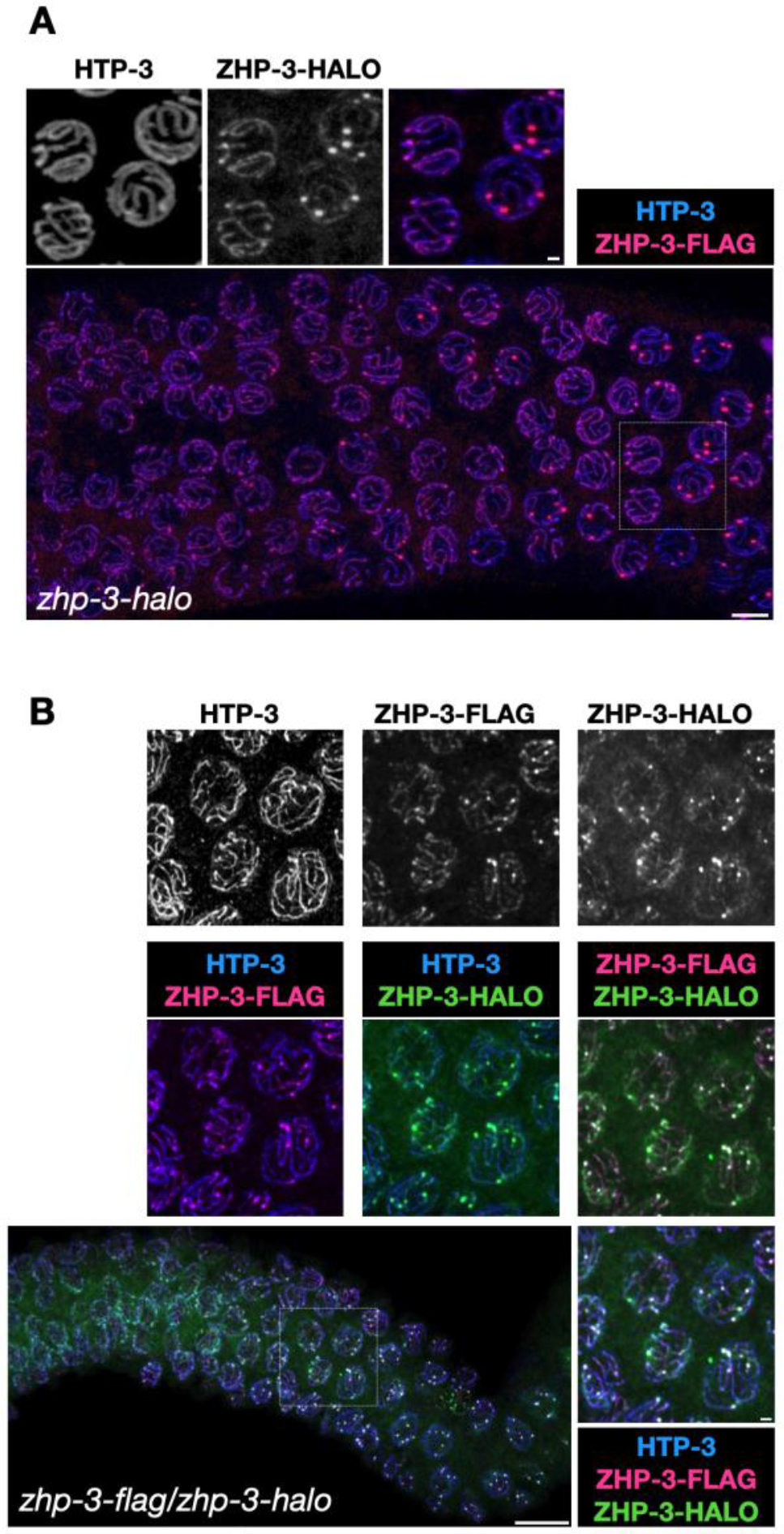
Cytological characterization of *zhp-3-halo* A. Confocal microscopy of HTP-3 and ZHP-3-FLAG in fixed *zhp-3-halo* homozygous worms (CA1387), showing the localization of ZHP-3-HALO before and after crossover formation. Meiosis progresses from left to right. Notably, in early pachytene ZHP-3-HALO localizes throughout the synaptonemal complex (marked by HTP-3) before forming distinct foci in late pachytene. B. Confocal microscopy of fixed *zhp-3::halo/zhp-3::flag trans-heterozygous* worms. Note the similar localization of the functional ZHP-3-FLAG and the sub-functional ZHP-3-HALO. Scale bars are 5 μm in the zoomed-out images and 1 μm in the zoom-in images.

**Figure S3:**
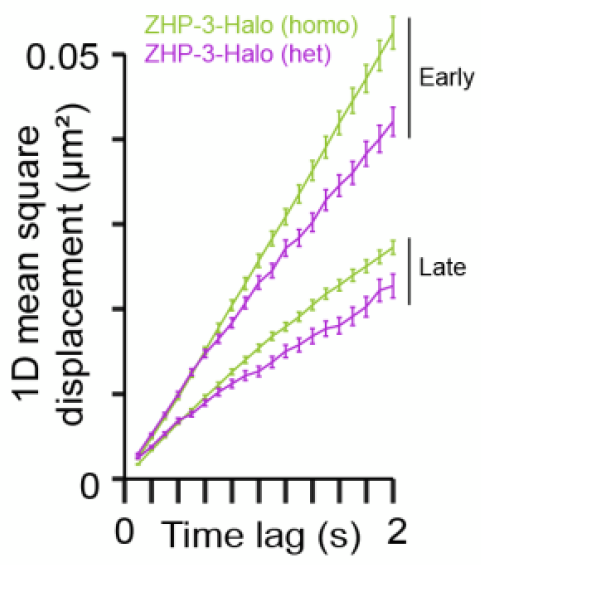
Single-molecule dynamics of the ZHP-3-Halo transgene in homozygote and heterozygote Comparison of mean square displacement curves acquired for ZHP-3-Halo molecules in the *zhp-3::halo/zhp-3::halo* (homo) and *zhp-3::halo/+* (het) genetic backgrounds.

**Figure S4:**
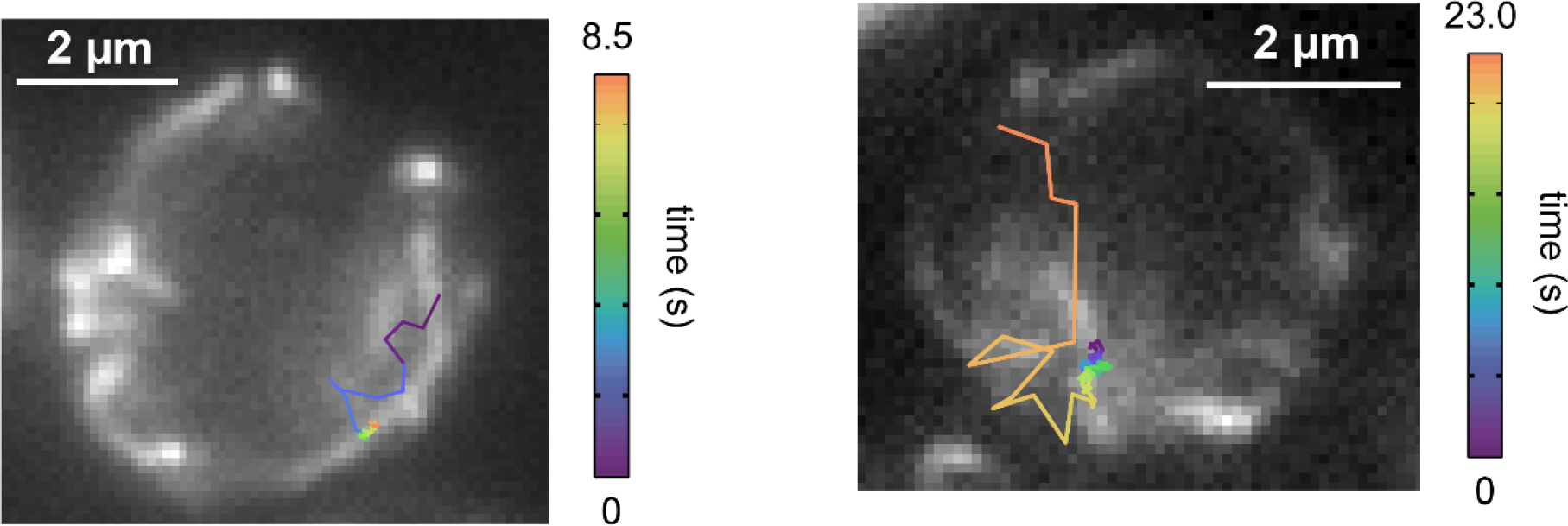
Example trajectories entering and leaving the SC-CR. Left: example trajectory of ZHP-3-Halo beginning in the nucleoplasm and ending in the SC-CR. Right: example trajectory of ZHP-3-Halo beginning in the SC-CR and ending in the nucleoplasm.

**Figure S5:**
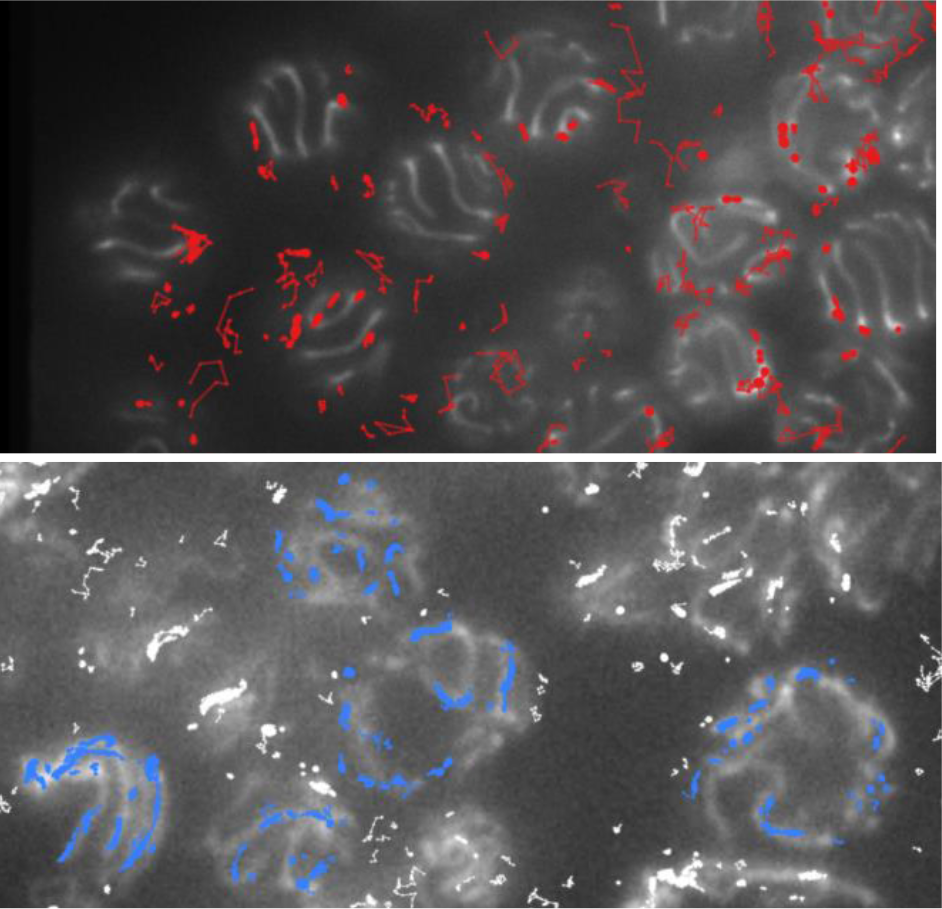
Comparison of labeling with PAJF549-HTL and JF552-HTL. Top: trajectories of ZHP-3-Halo labeled with PAJF549-HTL overlaid onto reference data of GFP-HIM-3. Bottom: trajectories of ZHP-3-Halo labeled with JF552-HTL overlaid onto reference data of GFP-HIM-3. Blue trajectories emphasize nuclei for which ZHP-3-Halo is clearly colocalized with stretches of the SC-CR.

**Figure S6:**
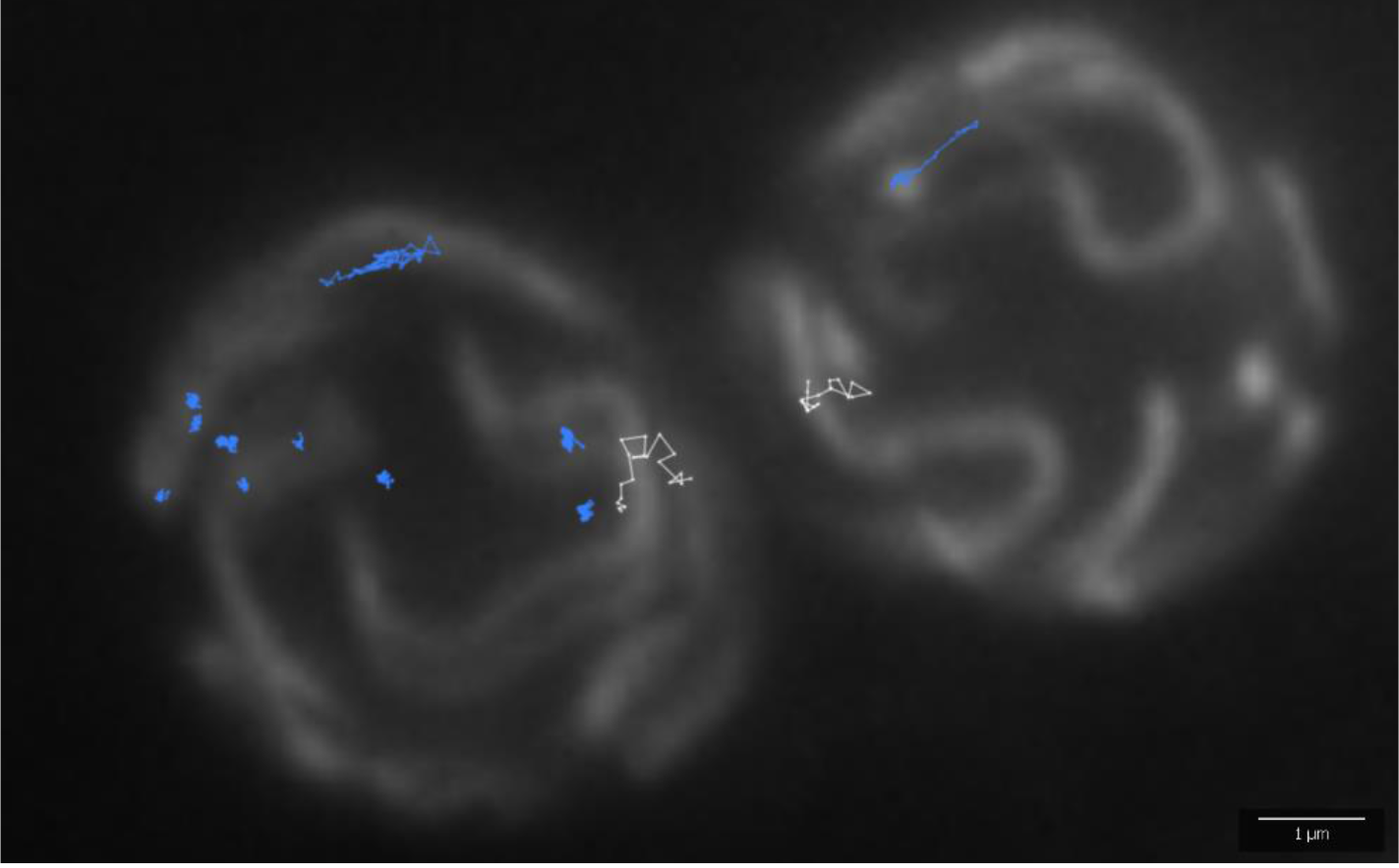
Example selected trajectories of ZHP-3-Halo. Blue: trajectories sufficiently associated with the SC-CR to be included for analysis. White: trajectories that are not clearly associated with the SC-CR and move too rapidly, and are not included.

## Notes

### Competing Interest Statement

The authors have declared no competing interest.

